# JQ1 Epigenetic Modulation of Pancreatic β-Cells (INS-1) Normalizes Glucose Sensitivity under Hyperglycemia: Therapeutic Preventive Implications for Type II Diabetes Mellitus

**DOI:** 10.1101/2023.08.15.553320

**Authors:** Mateo H. Hernández, Nazlı Uçar, Jude T. Deeney

**Affiliations:** John B. Alexander High School, Magnet for Health Sciences, 3600 E del Mar Blvd, Laredo, TX 78041; Department of Medicine, Evans Biomedical Research Center, Boston University Chobanian & Avedisian School of Medicine, 650 Albany Street, Boston, MA 02118

**Author notes:** [ ].

## Abstract

Chronic hyperinsulinemia and insulin resistance are prequels to type II diabetes mellitus (T2D), a disease that affects over 10% of Americans and is closely associated with obesity, as nearly 90% of diabetic patients are overweight or obese. Chronic excess nutrient exposure leads to glucolipotoxicity in pancreatic β-cells, which is characterized by a left shift in glucose concentration-dependent insulin secretion. Previous studies from our laboratory have demonstrated that the Bromodomain and Extra-Terminal (BET) protein inhibitor JQ1 (400 nM) increases fatty acid (FA) oxidation in clonal pancreatic β-cells (INS-1), leading to a partial reversal of glucolipotoxicity. The present research investigates the effect of JQ1 on the glucose sensitivity of INS-1 cells under hyperglycemic conditions. INS-1 cells were pre-treated with dimethyl sulfoxide-based 400 nM JQ1 for 3 days and cultured for 5 days in RPMI 1640 media (11 mM glucose). Subsequently, INS-1 cells were pre-incubated in a modified Krebs-Henseleit buffer (1 mM glucose) before adding test solutions (1-12 mM glucose). Samples for insulin release and cellular content were measured using a Homogeneous Time-Resolved Fluorescence insulin assay kit. Insulin secretion from INS-1 cells cultured in 11 mM glucose was maximally stimulated at 4 mM glucose. Treatment with JQ1 for one day right-shifted maximal glucose-stimulated insulin secretion (GSIS) to 6 mM glucose compared to control but did not prevent stimulated insulin secretion at 4 mM glucose. Three-day treatment with JQ1 reduced secretion at 4 mM glucose to basal level (1 mM) while maintaining the right-shifted concentration-dependent GSIS, an effect described herein for the first time. Total insulin content (TIC) and release as its percentage were also measured, indicating a higher TIC and lower percentage use in JQ1-treated cells. Additionally, lipid concentration was photographically fluorescent analyzed, showing significant depletion of lipid droplets in JQ1-treated cells. Results imply that epigenetic modulation of pancreatic β-cells with JQ1 beneficially alters signal transduction pathways that maintain insulin-glucose homeostasis by ameliorating glucolipotoxicity through the preservation of a low GSIS basal level, increment of GSIS maximum capacity, delay of GSIS cuspid level at 12 mM glucose, more efficient spend of total insulin content resources, and diminished lipid accumulation due to increased FA oxidation—thereby suggesting JQ1 returning hyperglycemic β-cells to physiological conditions. Suggested future directions include improving the efficacy and accuracy of JQ1 and similar small-molecule BET inhibitors through tests in human subjects’ isolated islets, as this presents an innovative course of research for the prevention of T2D and comorbidities.

## Introduction

Simultaneous persistently elevated levels of glucose in blood and insulin resistance precede the onset of type II diabetes mellitus (T2D), a disease that affects over 10% of Americans. Considering that nearly 90% of individuals diagnosed with diabetes are overweight or obese, T2D is closely associated with obesity. The continuous exposure to excessive nutrients over time culminates in a phenomenon known as glucolipotoxicity within pancreatic β-cells, the releasers of insulin. This state is distinguished by a shift towards the left in the concentration-dependent insulin secretion response to glucose—a regulatory process largely dictated by transcriptional mechanisms, wherein the bromodomain has emerged as a pertinent factor with metabolic implications.^**1**^

The exciting development of small molecule inhibitors of Bromodomain and Extra Terminal (BET) protein binding to chromatin has revealed that histone-protein interactions can be beneficially drugged. These inhibitors compete for the bromodomain’s acetyl-lysine binding pocket, thereby displacing the BET protein from chromatin, which alters the transcriptional activity of the target gene.^**2**^

Recently, particular attention has been centered on the activity of BET proteins as transcriptional co-activators for cell cycle and proliferation genes in oncological contexts. Instead, we further our investigation on the concept of BET proteins as co-repressors of transcription depending on the signal transduction, cellular, and gene context.

Interesting therapeutic opportunities may lie with small molecule inhibition of the co-repressor functions of BET proteins. These new developments have prompted a shift in attention beyond chromatin regulator function in malignancy to a broader concern that includes metabolism.^**3**^

A notably promising inhibitor, JQ1, was first synthesized in 2010 by the James Bradner Laboratory at Brigham and Women’s Hospital of Harvard Medical School and named after Lead Chemist Jun Qi, Ph.D..^**4**^ Since the beginning of the previous decade, exciting advancements have been made in the comprehension of the nature and action of this novel compound.

Previous studies from our laboratory have demonstrated that the BET protein inhibitor JQ1 at high concentrations (400 nM) increases fatty acid (FA) oxidation in clonal pancreatic β-cells (INS-1), leading to a partial reversal of glucolipotoxicity.^**2**^

The present research investigates the effect of JQ1 on the glucose sensitivity of INS-1 cells under hyperglycemic conditions. We investigate whether BET protein inhibition confers metabolic protection that may prevent the development of T2D.

## Methods

INS-1 cells were pre-treated with dimethyl sulfoxide (DMSO)-based active (+) and non-active (−) JQ1 compound (400 nM) for 1 and 3 days prior to measurement of insulin secretion and content.

INS-1 cells were pre-incubated at 37°C in RPMI 1640 media (11 mM glucose) for 5 days. Afterward, INS-1 cells were incubated at 37°C in 0.05% bovine serum albumin (BSA) Krebs-Henseleit (KREBS) solution buffer for 2 periods of 30 min, changing and replenishing the media in-between. Then, INS-1 cells were incubated at 37°C in different glucose concentrations of 1 mM glucose to 12 mM glucose in 0.05% BSA-KREBS solution buffer for 60 min.

Subsequently, specific antibodies were added to samples for measurement of insulin release and cellular content by a homogeneous time-resolved fluorescence (HTRF) insulin assay kit and incubated overnight. Using a TECAN M1000 Pro Plate Reader, the fluorescence-resonance energy transfer (FRET) signal between two separately labeled insulin-binding antibodies was measured.

Additionally, INS-1 cells were stained with green BODIPY (1 μg/ml) for 20 min at 37°C in RPMI media without serum. After stain removal, cells were imaged using a Nikon TE200 fluorescence microscope (20X magnification) equipped with a cooled Olympus DP72 CCD camera and Cellsense imaging software.

## Results

As an objective of our experiments, we obtained results within measurements of three factors relevant to the study of glucotoxicity, which characterizes the diabetic conditions investigated herein: (i) glucose-stimulated insulin secretion (GSIS), (ii) total insulin content (TIC) and percentage of its constant release, (iii) and lipid accumulation. In all reports, control represents that sample that was not treated with JQ1 at any time, thereby emulating the physiological conditions of hyperglycemia.

> (Note: In order to provide adequate space for displaying the figures of Results, the following three pages will be dedicated exclusively to one set of figures each. In *6*, it will be reserved for *Fig. 4* & *Fig. 5*; in *7*, it will be reserved for *Fig. 6* & *Fig. 7*; and in *8*, it will be reserved for *Fig. 8*.)

**Fig. 1.**
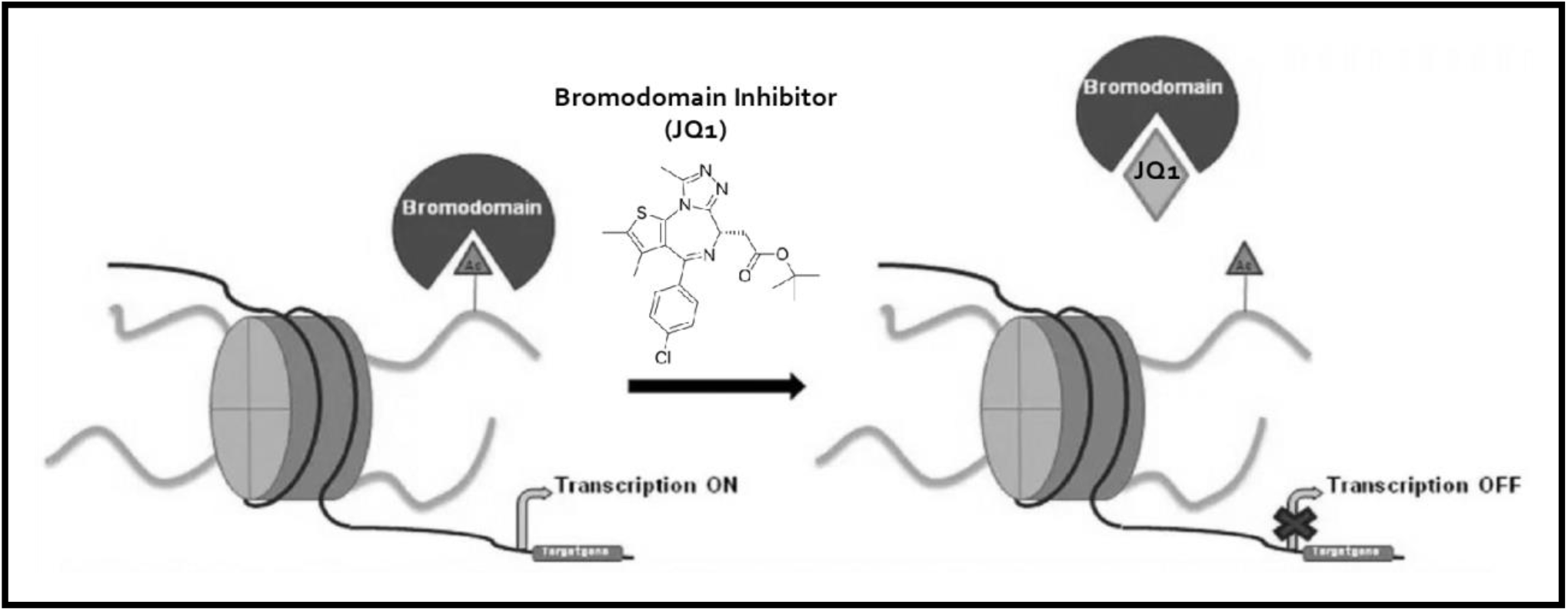
Molecular approach of JQ1 inhibiting bromodomain (BET)—found in epigenetic factors that control transcription. The bromodomain is a 110 amino acid motif comprised of four anti-parallel α-helices with two connecting loops that form a binding pocket for ε-acetyl-lysines of histones present in nucleosomal chromatin. *(Adapted from* Pérez-Salvia and Esteller^***6***^)

**Fig. 2.**
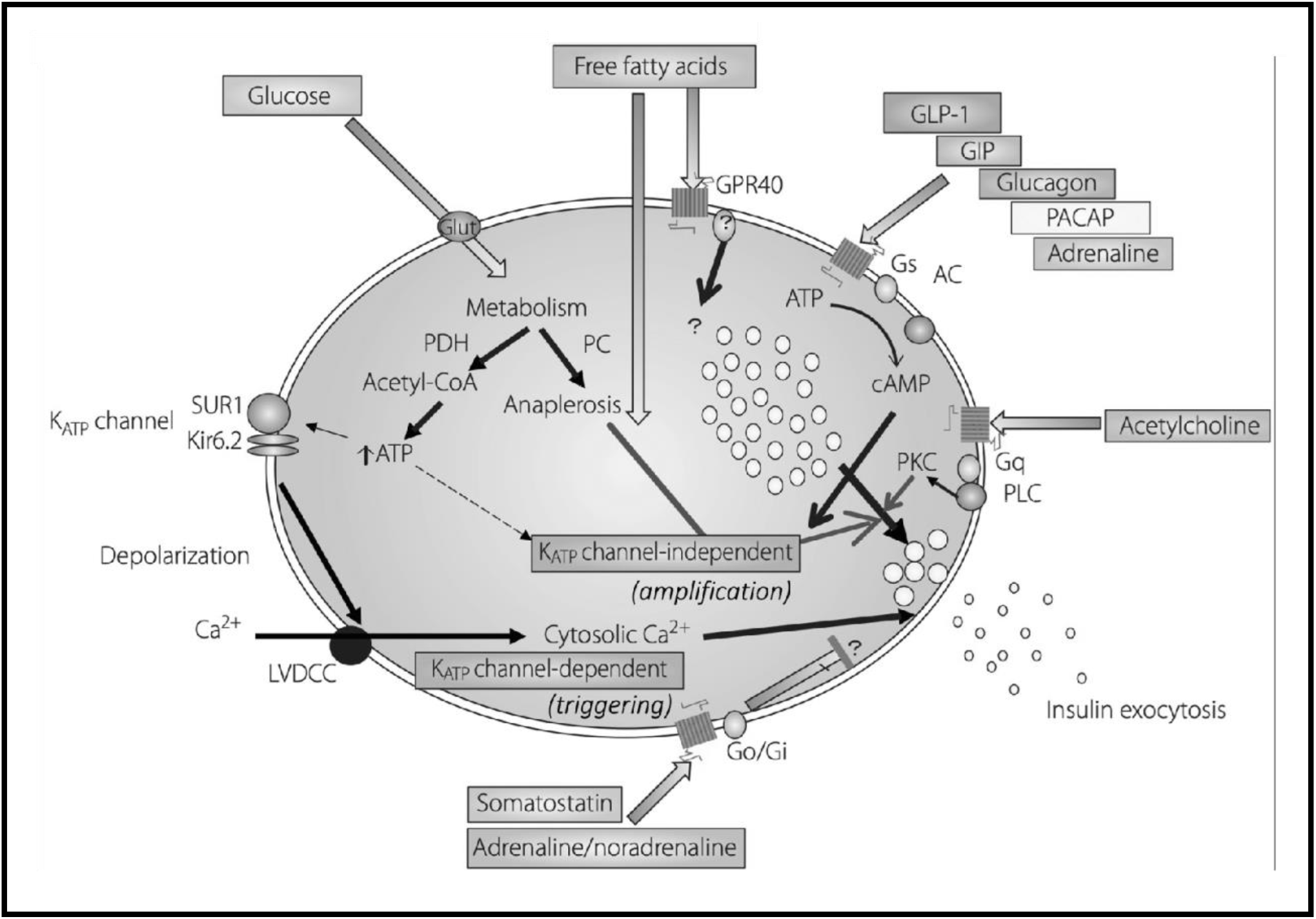
Physiological signaling Mechanism of GSIS in Pancreatic β-Cells. Glucose enters the pancreatic β-cell via the membrane glucose transporter, is phosphorylated by glucokinase, and undergoes glycolysis to form pyruvate. After its metabolization, pyruvate undergoes cellular respiration in the mitochondria, amplifying ATP concentration, which closes ATP-sensitive K^+^ channels, causing membrane depolarization that opens Ca^2+^ channels, increasing Ca^2+^ levels and thereby triggering insulin exocytosis. *(Adapted from* Komatsu et al.^***5***^)

**Fig. 3.**
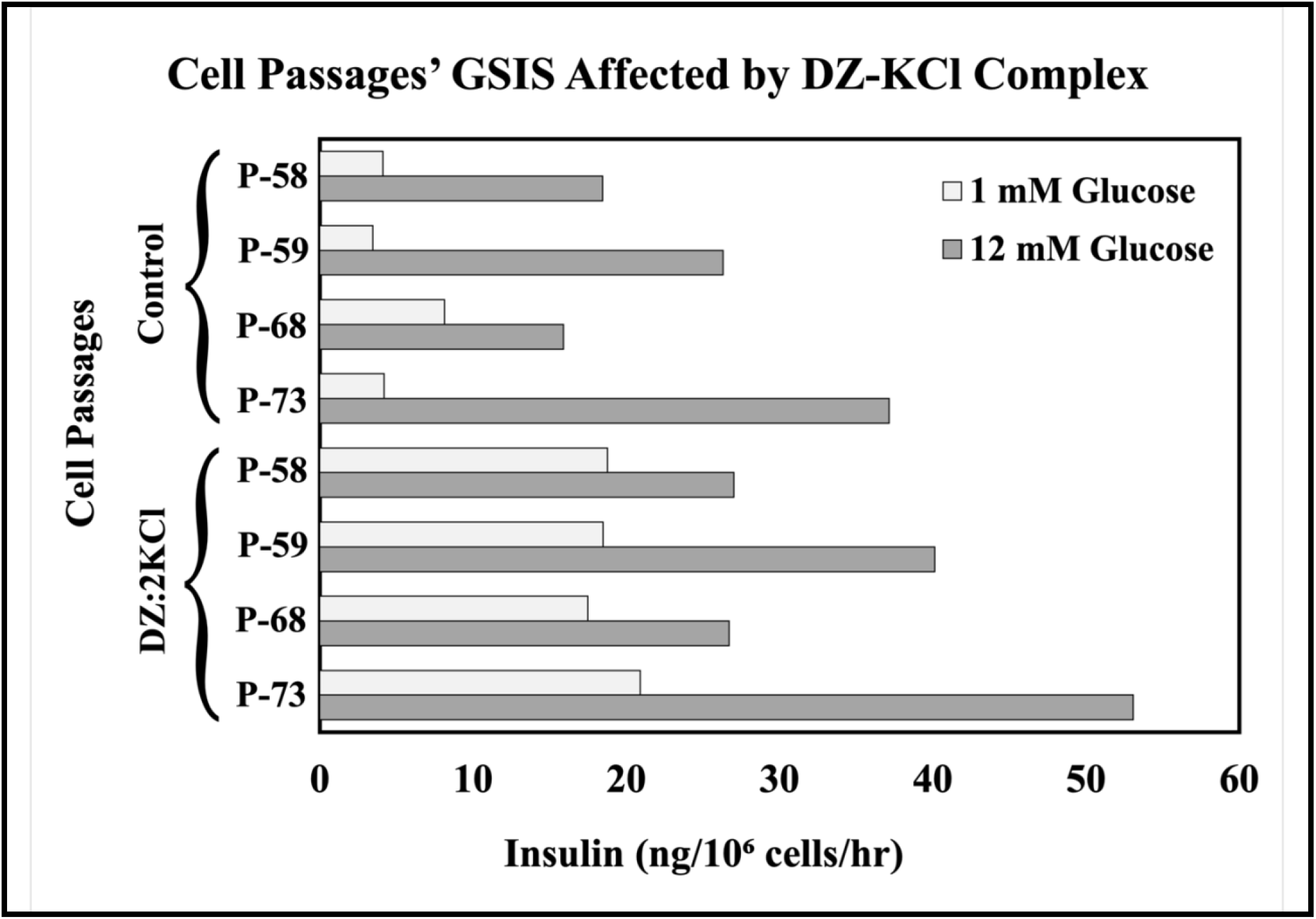
Selection of Cell Passage. INS-1 cell passage for the experiments was selected by testing the efficacy of four samples of P-58 through P-73 in GSIS directly affected by diazoxide (DZ) counteracted by potassium chloride (KCl)—P-73 was ultimately selected for the following experiments.

**Fig. 4.**
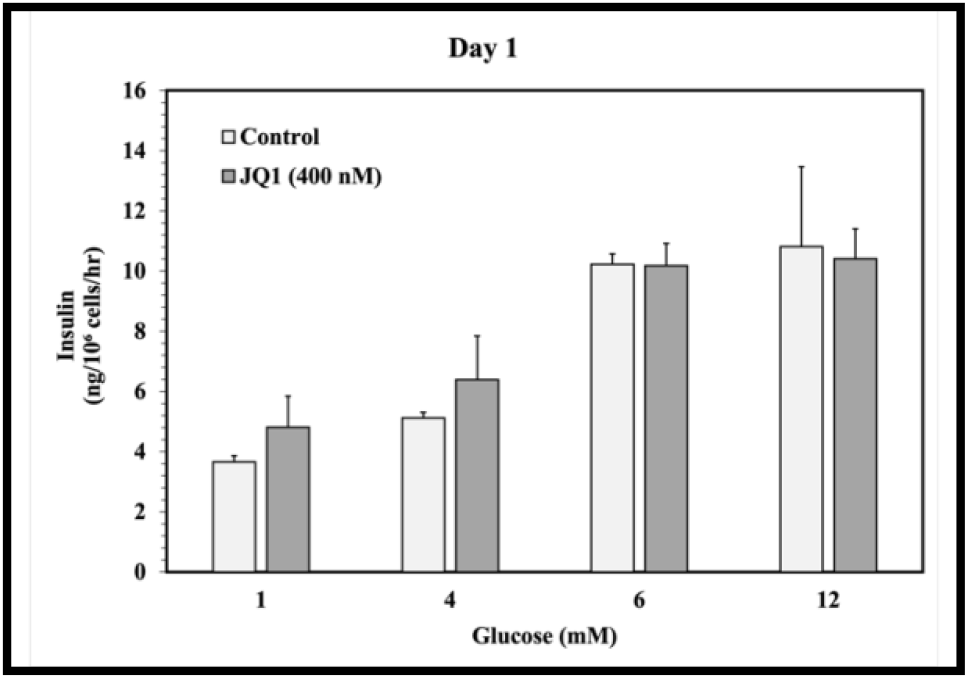
GSIS of INS-1 cells, 1 day post JQ1 treatment. After the first day, control basal GSIS levels comprehended up to 1 mM glucose, increasing from 4 mM glucose and reaching maximum capacity at 6 mM glucose. Similarly, JQ1-treated cells GSIS had relatively analogous behavior and maximum.

**Fig. 5.**
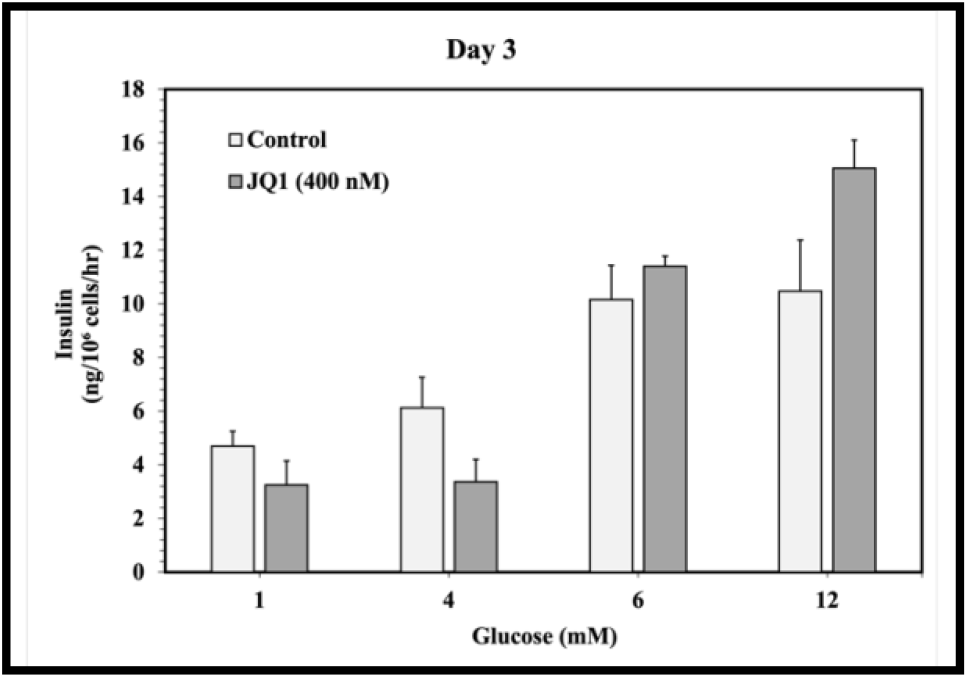
GSIS of INS-1 cells, 3 days post JQ1 treatment. After the third day, control GSIS had similar behavior to that of the first day. Contrastively, JQ1-treated cells basal GSIS levels comprised up to 4 mM glucose, increasing from 6 mM glucose and reaching maximum capacity at 12 mM glucose.

**Fig. 6.**
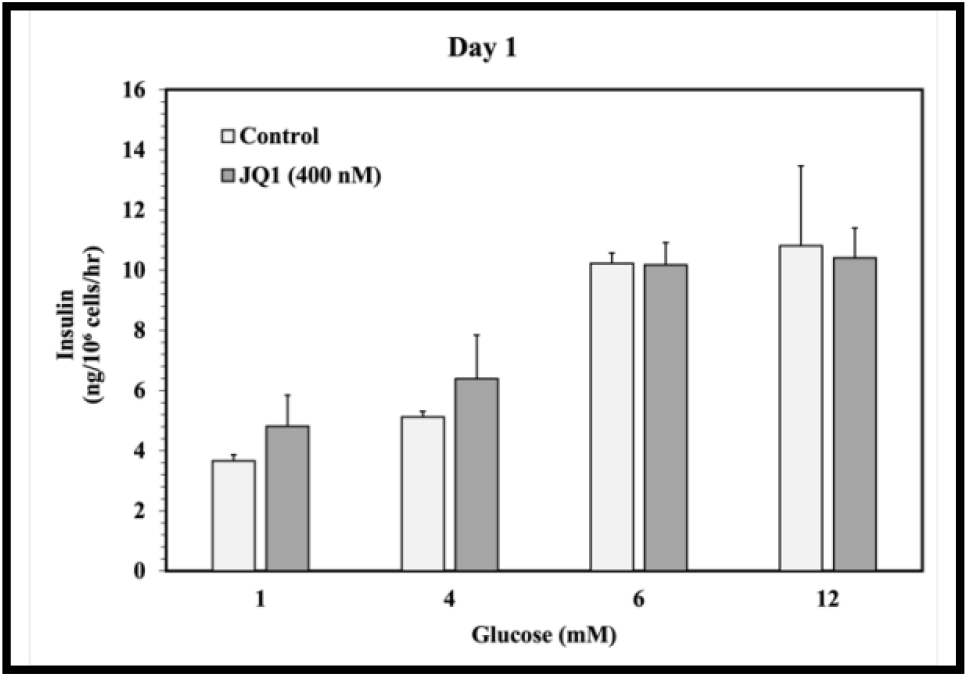
Relative constant TIC remained greater under JQ1 treatment (from Day 3). Finalizing, control TIC levels were lower—by ∼33%—than those of JQ1-treated cells. This result demonstrates that control used a significantly higher percentage of its insulin reserves per cell in GSIS in all instances. Furthermore, JQ1-treated cells used a lesser percentage of reserves while secreting even more than control throughout the major glucose levels.

**Fig. 7.**
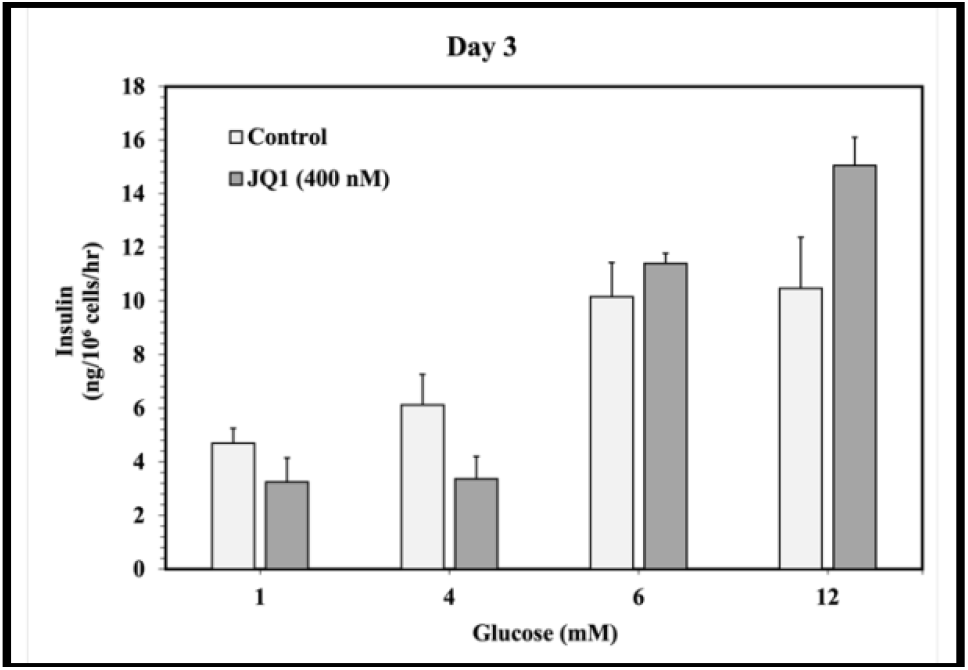
Percentage of TIC released hourly by INS-1 cells, 3 days post JQ1 treatment. After the third day, control basal levels of TIC released hourly comprehended up to 1mM glucose, increasing from 4mM glucose and reaching maximum capacity at 6mM glucose. Contrastively, JQ1-treated cells basal levels of TIC released hourly comprehended up to 4 mM glucose, increasing from 6 mM glucose and reaching maximum capacity at 12 mM glucose.

**Fig. 8.**
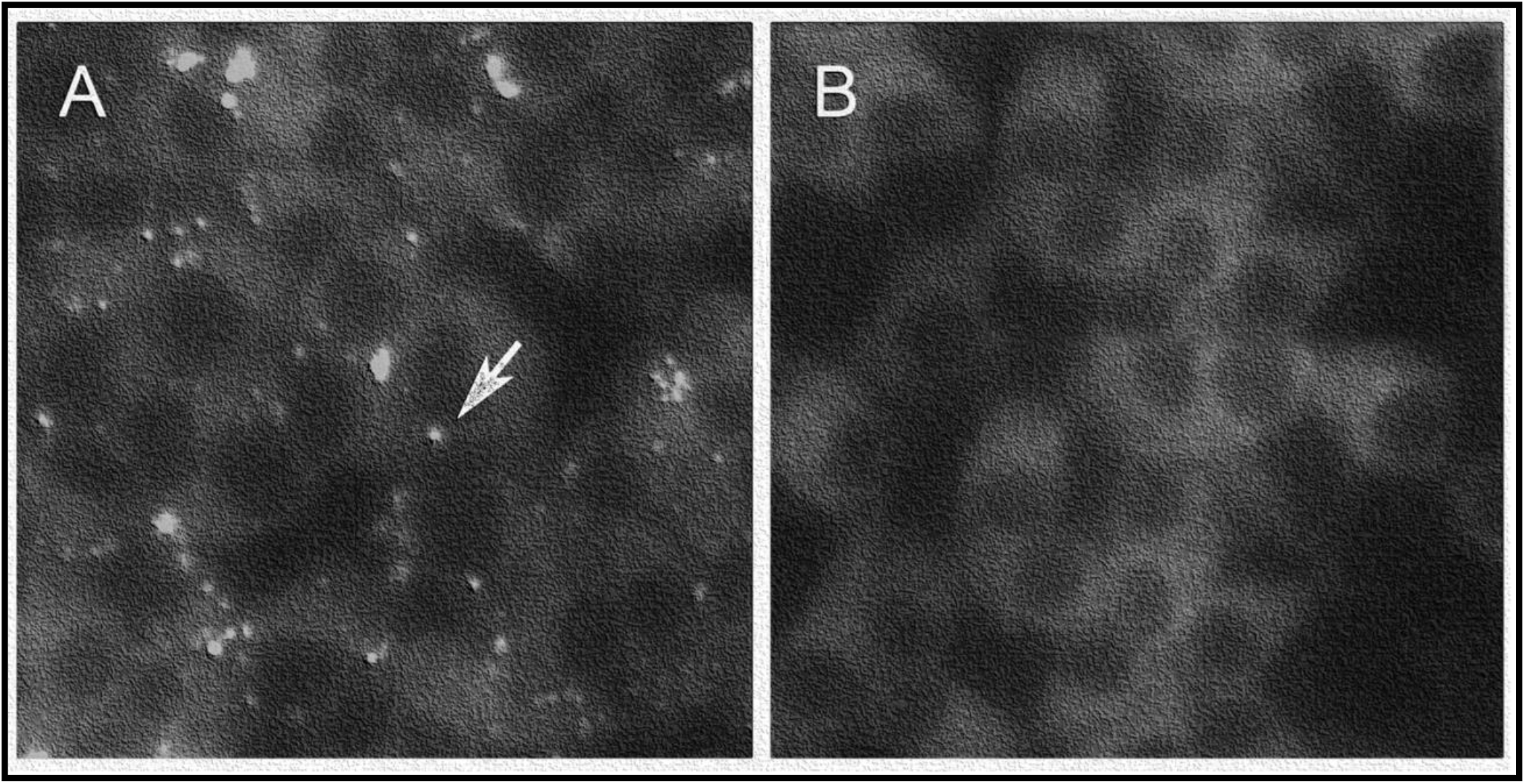
JQ1 reduces lipid accumulation in INS-1 cells cultured in 11 mM glucose. *A* shows non-treated control cells; *B* shows cells with JQ1 after three days. The white arrow points to a lipid droplet in *A*.

i. The GSIS of the INS-1 cells was measured in two instances—24 and 72 hours after JQ1 (400 nM) application—in nanograms per million cells per hour.
ii. The TIC of the INS-1 cells was measured at the end of the experiments—72 hours after JQ1 (400 nM) application—in nanograms per million cells.
iii. The lipid accumulation of the INS-1 cells was measured at the end of the experiments—72 hours after JQ1 (400 nM) application—by means of photographic analysis.

## Discussion

We have demonstrated multiple beneficial effects of BET protein inhibition on β-cell function in our experiments. The first benefit is the potentiation of insulin transcription, resulting in elevated insulin secretion in INS-1 cells as shown in *Fig. 4* & *5*. Here, we are the first source to describe the *Right-Shift Effect* of JQ1 upon concentration-dependent GSIS—as referenced in *Fig. 9*.

**Fig. 9.**
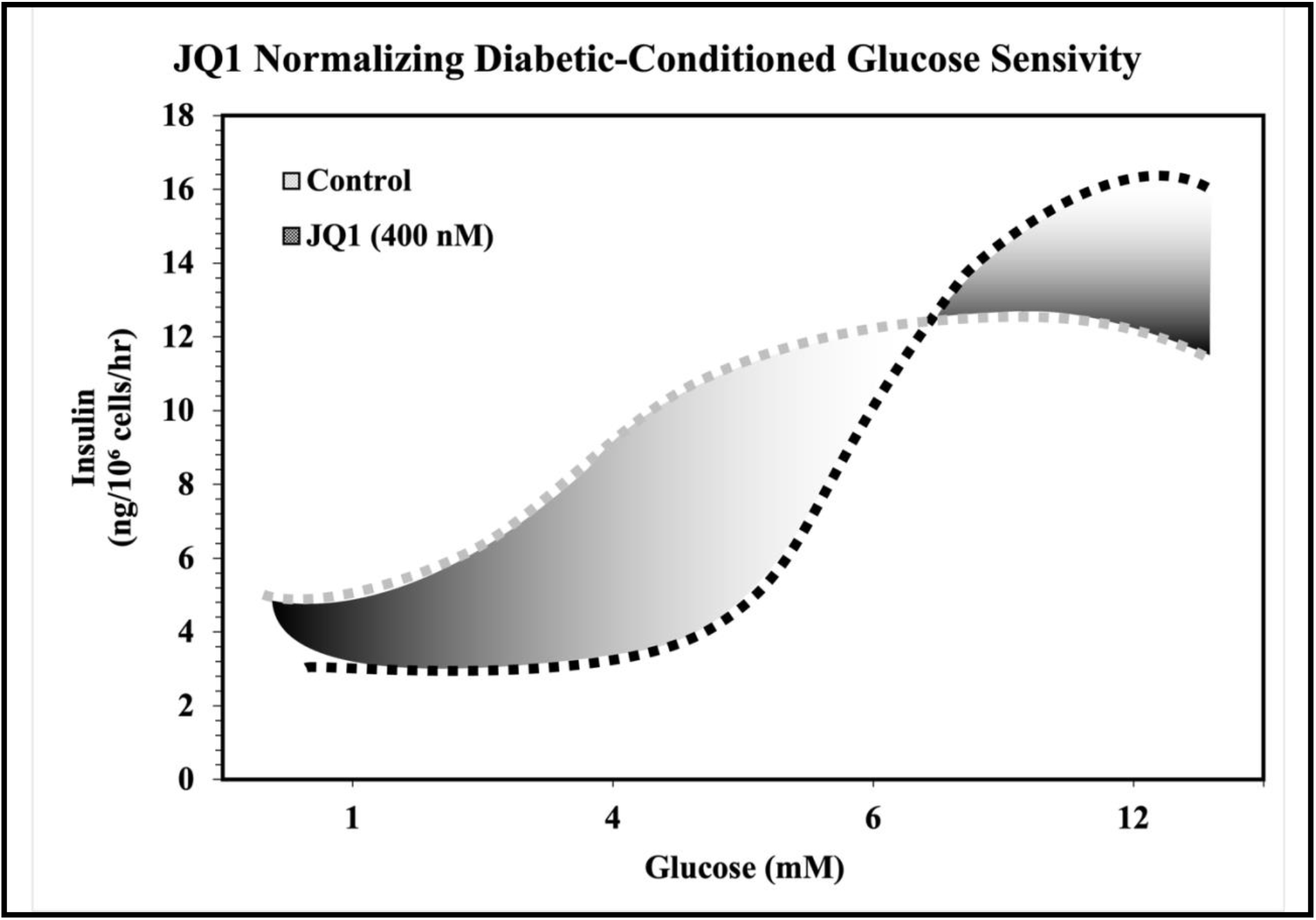
The “Right-Shift Effect” of JQ1 upon concentration-dependent GSIS (from Day 3). In JQ1-treated cells, GSIS basal levels remained lower than control (< 4 ng/10^6^ cells/hr insulin), GSIS maximum capacity was delayed, and GSIS cuspid level at 12 mM glucose was incremented by >30% compared to control. This exciting result indicates the reversal of the main factor of glucolipotoxicity and the return to physiological, non-diabetic conditions in INS-1 cells.

The second benefit lies in the observation that INS-1 cells cultured with JQ1 under hyperglycemic conditions (11 mM glucose) exhibit increased insulin reservoirs while releasing more insulin than typical β-cells as shown in *Fig. 6*. Hence, the modest quantity of released insulin as a percentage of TIC, as shown in *Fig. 7*, —coupled with the favorable impact of diminishing surplus intracellular lipid on insulin mRNA^**3**^—could collectively contribute to the clear fold-increase in insulin content in INS-1 cells compared to normal β-cells after BET protein inhibition with JQ1.

The third benefit is a notable alteration in β-cell fuel utilization, resulting in increased FA oxidation and decreased intracellular triglyceride stores as shown in *Fig. 8*. Ordinarily, glucose acts to inhibit FA oxidation through anaplerosis within the citric acid cycle and associated downstream reactions. However, when these fuels are chronically elevated, as is the case in obesity and T2D, β-cells accumulate excess lipid that eventually impairs function, including insulin release.^**2**^ Inhibition of BET proteins dampens the inhibitory effect of glucose upon FA oxidation, thus permitting its increased rates and preventing excess lipid accumulation—thereby potentially providing protection from glucolipotoxicity.

## Conclusions

Our results imply that epigenetic modulation of pancreatic β-cells with JQ1 beneficially alters signal transduction pathways that maintain insulin-glucose homeostasis by ameliorating glucolipotoxicity to enhance glucose sensitivity and reverse insulin resistance.

Exposure to JQ1 led to an augmentation in insulin secretion within INS-1 cells— suggesting potential therapeutic benefits for humans. The reduction in intracellular lipid content could arise from various altered metabolic pathways, encompassing heightened lipolysis, diminished triglyceride synthesis, or amplified FA oxidation—significantly contributing to the progressive reduction of intracellular lipid droplets.

By implementing molecular BET inhibitors, patients with insulin resistance might benefit from both increased insulin stores and enhanced β-cell capacity to oxidize FA—protecting the β-cell from excess nutrient-induced damage (glucolipotoxicity). Thus, the possibility that improved, chromatin-directed inhibitors such as JQ1 could offer benefits for obese or diabetic patients represents an innovative and noteworthy avenue, warranting further comprehensive investigation.

Epigenetically altered metabolic pathways found in the β-cell could be replicated in other tissues, such as myocytic, adipose, and hepatic, to provide a multi-organ defense that impedes the onset or mitigates the severity of T2D and comorbidities through the reversal of glucolipotoxicity.

## Acknowledgments

I express my utmost gratitude to Dr. Jude T. Deeney for his invaluable guidance in this project. Furthermore, I sincerely appreciate the pivotal support and insight that Ms. Nazlı Uçar provided me throughout the entirety of the RISE Program—an opportunity for which I am grateful to Boston University. I am also thankful for the memorable experience lived with my lab partners, Alexander C. Yeung and Annabelle S. Chung, during this summer.

We thank the Boston University Chobanian & Avedisian School of Medicine Analytical Instrumentation Core, directed by Dr. Lingyi L. Deng and co-directed by Dr. Matthew B. Au, for providing the infrastructure for this project.

